# The greening-causing agent alters the behavioral and electrophysiological responses of the Asian citrus psyllid to a putative sex pheromone

**DOI:** 10.1101/2023.11.09.566442

**Authors:** Haroldo X. L. Volpe, Michele Carmo-Sousa, Rejane A. G. Luvizotto, Renato de Freitas, Victoria Esperança, Josiane C. Darolt, Abner A. L. Pegoraro, Diego M. Magalhães, Arodi P. Favaris, Nelson A. Wulff, Marcelo P. Miranda, José Maurício S. Bento, Walter S. Leal

## Abstract

The Asian Citrus Psyllid (ACP), *Diaphorina citri,* is a vector of the pathological bacterium *Candidatus* Liberibacter asiaticus (CLas), which causes the most devastating disease to the citrus industry worldwide, known as greening or huanglongbing (HLB). Earlier field tests with an acetic acid-based lure in greening-free, ‘Valencia’ citrus orange groves in California showed promising results. The same type of lures tested in São Paulo, Brazil, showed unsettling results. During the unsuccessful trials, we noticed a relatively large proportion of females in the field, ultimately leading us to test field-collected males and females for *Wolbachia* and CLas. The results showed high rates of *Wolbachia* and CLas infection in field populations. We then compared the olfactory responses of laboratory-raised, CLas-free, and CLas-infected males to acetic acid. As previously reported, CLas-uninfected males responded to acetic acid at 1 µg. Surprisingly, CLas-infected males required 50x higher doses of the putative sex pheromone, thus explaining the failure to capture CLas-infected males in the field. CLas infection was also manifested in electrophysiological responses. Electroantennogram responses from CLas-infected ACP males were significantly higher than those obtained with uninfected males. To the best of our knowledge, this is the first report of a pathogen infection affecting a vector’s response to a sex attractant.

## Introduction

The Asian Citrus Psyllid (ACP), *Diaphorina citri* (Hemiptera: Psyllidae), is a vector of the bacterium *Candidatus* Liberibacter asiaticus (CLas), which causes the citrus disease known as greening or huanglongbing (HLB)^1^. This pathogen parasitizes the phloem and blocks nutrient circulation, thus causing citrus trees to become unproductive^1^. Citrus growers suffer severe losses manifested in reduced fruit quality and production and, ultimately, plant death.^2^ It has been demonstrated that CLas induces infected plants to produce an ACP attractant, methyl salicylate^3^, leading to ACP’s preference for CLas-infected over healthy plants. ACP feeds on infected plants, but due to diseased plants’ low nutritional value, the psyllids move to healthy plants to complete feeding, thus vectoring the HLB-causing agent^3^ from infected to healthy trees. Citrus growers eradicate infected trees to decrease the bacterium’s spread and, consequently, sustain severe losses in production.

Florida used to be the largest citrus producer in the United States of America for many years, with more than 300 million boxes produced in 1997-1998 and continuous yearly production of more than 250 million boxes from 1992-1993 until 2003-2004^4^. After HLB detection in the state in 2005^5^, the Florida citrus industry has been declining precipitously from 169.25 million boxes in 2004-2005^4^ to 15.85 million in 2022-2023, as of July 12, 2023^6^, i.e., suffering a 90.6% reduction. As Florida’s citrus industry is being decimated, California, with greening-free commercial orchards, became the nation’s largest producer since 2016-2017. Last year, California and Florida contributed 61.8% and 36% of the nation’s citrus production, respectively^4^. By contrast, Florida contributed 79.7% and California 17.4% of the nation’s output in 2003-2004^4^ before HLB infestation in Florida^5^. The vector and the bacterium have been detected in California, but strict quarantine measures have prevented the bacterium from reaching commercial citrus plants. More than 5,000 HLB-infected trees have been found and removed from residential areas^7^. At the time of this writing, there are reports of infected ACP nymphs being detected in commercial orchards^7^.

Brazil, the largest orange producer in the world^8^, has been sustaining severe losses since 2004^9^ when greening was discovered for the first time in the State of São Paulo^10–12^. Because there are no cost-effective treatments for infected plants^13^, 61,585 ha have been eradicated from 2018 to 2021 to contain the spread of greening. Although eradication was alleviated with renewed and expanded areas, the productive acreage was reduced by 3.56%^9^. In the last three years, orange growers had to eradicate 7.26, 7.65, and 6.68% of the planted areas^14^.

The most effective approach to slow the spread of the diseases is vector control combined with scouting and replacing infected trees with HLB-free trees produced in screened vector-free nurseries. Vector control relies on applications of insecticides, particularly those with active ingredients like biphentrin, imidacloprid, and malathion^15^. However, alternative approaches are sorely needed, given the high cost of multiple insecticide applications, psyllid resistance, adverse effects on non-target species and beneficial insects, and impact on human health^13^. Chemical ecology-based approaches employing ACP attractants and/or repellents may contribute to the integrated management of this vector. Whereas repellents could be used in push-and-pull strategies^16^, pheromones, and other attractants may be employed for monitoring, surveillance, and possibly control strategies, such as attract-and-kill^17^ and mating disruption^18, 19^. Indeed, many chemical ecology-based efforts are currently being explored (review in ^13^). Previously, we have identified acetic acid as a sex attractant^20^, which we labeled a “putative” sex pheromone, given the analytical challenge to determine unambiguously whether this semiochemical is released only by ACP females. Field tests in HLB-free groves in California suggested that acetic acid significantly increased trap captures at specific doses, thus suggesting its potential in monitoring ACP populations^21^. As reported here, subsequent field tests in Brazil were disappointing. Captures in traps baited with acetic acid did not significantly differ from those in control traps. We surveyed ACP populations in the test area and found that CLas infected more than 98% of males and females. We then raised CLas-free and CLas+ colonies and compared males’ behavioral responses to acetic acid. We recapitulated the earlier findings with CLas-free males showing strong responses to acetic acid.^20^ At the same dose, CLas+ males did not significantly prefer acetic acid. However, at 50x higher doses CLas+ males were attracted considerably to acetic acid. Interestingly, acetic acid elicited significantly higher electroantennographic responses from CLas+ than CLas-free males. In summary, we report here that HLB infections alter ACP males’ behavioral and electrophysiological response to a sex pheromone.

## Results and Discussion

### Inconsistent Asian Citrus Psyllid captures in acetic acid baited traps

Previously, we have tested a slow-release formulation (ChemTica-A) of the Asian Citrus Psyllid (ACP) putative sex pheromone, acetic acid^20^, in an unsprayed ‘Valencia’ citrus orange grove at the California State Polytechnic University at Pomona, CA, with promising results^21^. Although the ACP density at the time of the experiments was very low, traps baited with ChemTica-A captured significantly more ACP males than control traps (N = 24, 1.87 ± 0.34 and 0.50 ± 0.10 males per trap per day in treatment and control traps, respectively; P=0.0001, Mann-Whitney test^21^). We failed to recapitulate this trap performance in 2019 in an area with natural ACP infestation in Mogi Mirim, State of São Paulo, Brazil (22° 25’ 55’’S; 46° 57’ 28’’W). Traps baited with ChemTica-A showed similar performance to control traps: 0.38 ± 0.06 and 0.42 ± 0.06 males per trap per day in treatment and control traps, respectively (N = 120, P = 0.6536, Mann-Whitney test).

To rule out possible formulation problems, we tested traps baited with a homemade formulation, which previously performed similarly to ChemTica-A in our field tests in Pomona^21^. This time, ACP captures in control and treated traps were not significantly different (0.96 ± 0.10 and 0.90 ± 0.10 males per trap per day in the control and homemade traps, respectively; N = 120; P = 0.7617, Mann-Whitney test). During these experiments in the State of São Paulo, we observed more females than males in adults captured in acetic acid and control traps. These findings prompted us to investigate whether the sex ratio of the ACP population in the experimental plot reflects the female biases in control and treatment traps.

### ACP sex ratio in an experimental organic plot

From February to December 2020, we collected 25 data points, each with 5 samples of 20 adults (100 insects per data point). The adult psyllids were aspirated from different plants in the experimental area. Two samples, collected on March 9 and November 27, showed a higher proportion of males, with male/female ratios of 1.10 ± 0.16 and 1.17 ± 0.14, respectively (Figure 1). Twenty-three samples showed a higher proportion of females than males, with male/female ratios ranging from as low as 0.3 to 0.93. The overall mean of 0.69 ± 0.04 suggests that throughout the 2020 flight season, on average, the field population comprised 43% more females than males (Figure 1).

**Figure 1.**
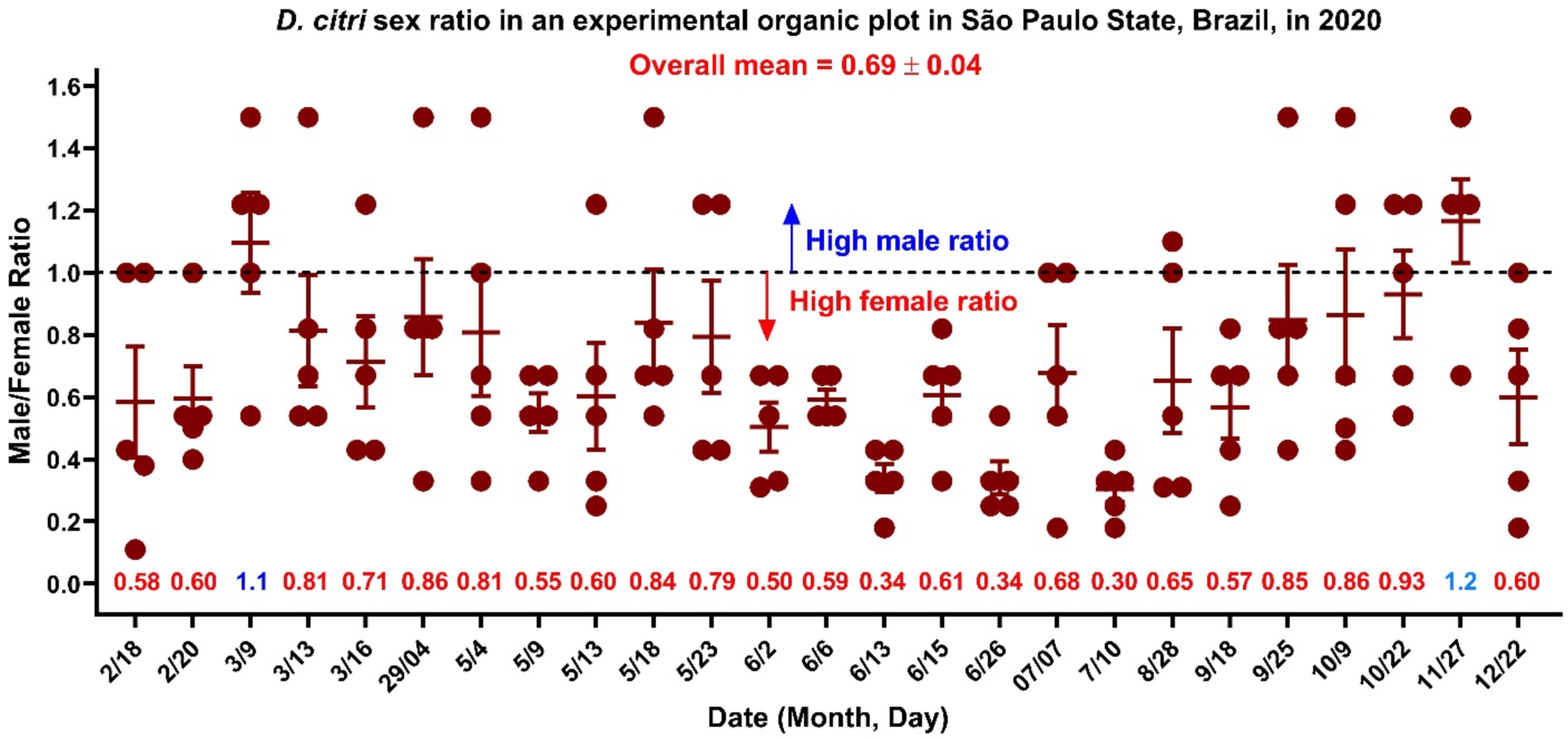
ACP male to female ratio in an experimental organic plot in Mogi Mirim, São Paulo State, Brazil, in 2020. Twenty-five data points were collected from February to December 2020 in Mogi Mirim. Twenty adults per plant were aspirated one by one from five plants sampled within the experimental area for each data point (of 100 adults). In the laboratory, adults were sexed, and the male/female ratios were recorded. The dotted line marks an equal number of males and females. Out of 125 samples (from a 25-day collection), only 20 showed a high male ratio (male/female ratio above 1). By contrast, 93 data points are below the dotted line, thus indicating a high female proportion. The overall mean suggests that the sex ratio was biased throughout the season toward females.

### *Wolbachia* infection rate in an experimental organic plot

Having observed a female bias in the ACP population in the experimental area in the State of São Paulo and the lack of response to the putative sex pheromone, we selected 1,100 out of the 2,500 collected adults to determine by the polymerase chain reaction (PCR) the level of infection with *Wolbachia* spp.^22^ and the *Candidatus* Liberibacter asiaticus (CLas)^23^ in this population. Of 1,100 samples, 1,084 (from 643 females and 441 males) were in good condition and analyzed by PCR. They represent 11 data points collected from February 18 to May 23, 2020 (Figure 2). All 410 males in samples collected in 10 data points were positive for *Wolbachia*. One of the 31 males collected on February 18, 2020, was negative; overall, one out of 441 males (0.24%) tested negative for *Wolbachia*. All 538 females representing nine data points were positive, whereas one of the 49 and one of the 56 females collected on March 9 and May 23, 2020, tested negative for *Wolbachia*. In short, we tested 643 females, and only two (0.31%) tested negative for *Wolbachia*. Although our data pertains to an experimental organic plot, it is known that *Wolbachia* is already fixed in the populations of *D. citri* distributed in agricultural settings in several states in Brazil, especially São Paulo^24^. Likewise, our laboratory colony derived from psyllids collected in Santa Fé do Sul, São Paulo (20° 12’ 43’’S; 50° 55’ 38’’W) and kept on healthy plants for almost 14 years showed high levels of *Wolbachia* infection. One hundred and two psyllids out of 103 analyzed insects (99.03%) tested positive for *Wolbachia* with a mean cycle threshold (C_t_) of 16.6 ± 0.1. Therefore, measuring behavior (e.g., attraction to semiochemicals) with our laboratory colony is more likely to emulate behavioral responses with wild-type psyllids from citrus groves in the State of São Paulo. Many factors, including dispersal ^25^, may contribute to psyllid sex ratios. It is conceivable that the skewed sex ratio observed in this experimental organic plot (Figure 1) was mediated, at least in part, by *Wolbachia* infection^26^.

**Figure 2.**
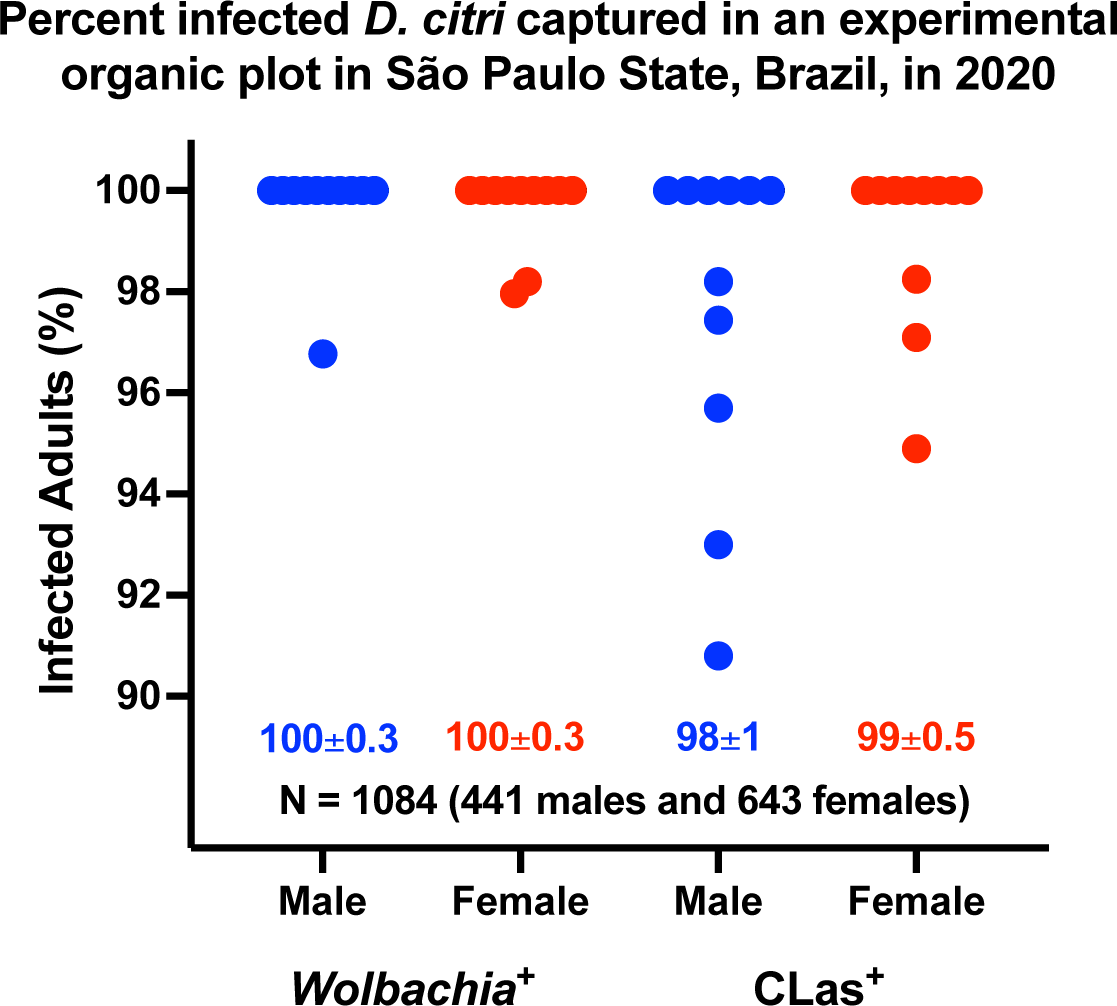
Percent of male and female ACP naturally infected with *Wolbachia* and CLas. Samples were collected for eleven days from February 18 to May 23, 2020, from an experimental organic plot in Mogi Mirim in the State of São Paulo, Brazil. Each point represents the percentage of adults infected with the tested bacteria. One thousand and one hundred psyllids were collected, but only 1,084 samples passed the quality test for PCR analysis.

### CLas infection rate in an experimental organic plot

CLas infection rate was also very high (Figure 2). Male samples from six data points were 100% positive. Only 1, 3, 3, 2, and 2 males were negative from 39, 43, 34, 47, and 42 males collected on March 16, April 29, May 12, 18, and 23, 2020, respectively. Female samples from eight data points were 100% positive for CLas. Female samples collected on March 13 (57 females), May 4 (59), and May 12 (65) had 1, 3, and 2 CLas-negative females, respectively. In summary, the level of CLas infection was surprisingly high in the experimental area. Although there are no data in the literature specific for Mogi Mirim (in the eastern part of São Paulo State), it has been reported that the percent of CLas-infected ACPs was constant throughout the year in the southwestern region of São Paulo State and, in average, 65.3%^27^. Data from 2014-2017 ranged from 33% in the northern area of São Paulo State/Triângulo Mineiro from Minas Gerais State to 74.6% in the southwestern part of the state^27^.

It is well documented in the literature that pathogen infections may cause changes in a vector’s behavior that ultimately lead to the enhanced transmission of the pathogen^28, 29^, a phenomenon known as the host manipulation hypothesis^30^. For example, *Plasmodium falciparum* – a protozoan that causes malaria in humans – increases the frequency of multiple feeding in its vector *Anopheles gambiae*^31^. Likewise, the tomato yellow leaf curl virus (TYLCV) - vectored by *Bemisia tabaci* – enhances the insect vector feeding behavior by increasing contacts and longer durations of salivation into phloem sieve elements^32^.

The *Candidatus* Liberibacter asiaticus is no exception. *Ca* Liberibacter spp. infection causes a change in metabolism^33–35^, fitness^36^, feeding^37^, and dispersal behavior^36^. It is worth mentioning that CLas infection may also shorten ACP lifespan,^38^ thus negatively affecting the transmission window. Here, we asked whether CLas infection could affect male responses to the putative female sex attractant^20^.

### Olfactory responses of CLas-free (CLas^-^) ACPs to acetic acid

Using a multiple-choice olfactometer^39^, we first measured the responses of uninfected ACP males to acetic acid^20, 21^. As reported above, our laboratory colony is naturally infected with *Wolbachia* (99.03%). Seven-day-old virgin males from a CLas-free laboratory colony, reared on orange jasmine, *Murraya paniculata*^20^, were tested under periodicity, luminosity, relative humidity, and temperature conditions to mimic the natural conditions in most citrus fields in Brazil^20^. Male responses were expressed as mean time spent in an odorant or control field. The arena had two control and two treatment fields. Psyllids were tested one at a time to allow accurate measurement of the time they spent on each of the two fields (one with acetic acid and the other with solvent control). Males who did not cross the odorant or control field lines within 5 min were recorded as “non-responders.” Otherwise, psyllids were observed for 10 min.

One hundred and ten ACP males were tested for their responses to acetic acid at 1 µg (100 µl of 0.01 µg/µl solution in hexane; 100 µl of the solvent for control) loaded on a cotton swab. Sixteen males did not respond. The responding males (N = 94) showed a significant preference for the odorant field (1 µg acetic acid; residence times: treatment 5.8 ± 0.3 min, control 4.2 ± 0.3 min; P = 0.0073, Wilcoxon matched-pairs signed rank test; hereafter referred to as Wilcoxon test) (Figure 3A). As a negative control, we tested 7-day-old virgin females. The female responders (N = 75, tested 90) showed no preference (residence time on the control field, 5.1 ± 0.4 min; treatment, 4.9 ± 0.4 min; P = 0.8510, Wilcoxon test).

**Figure 3.**
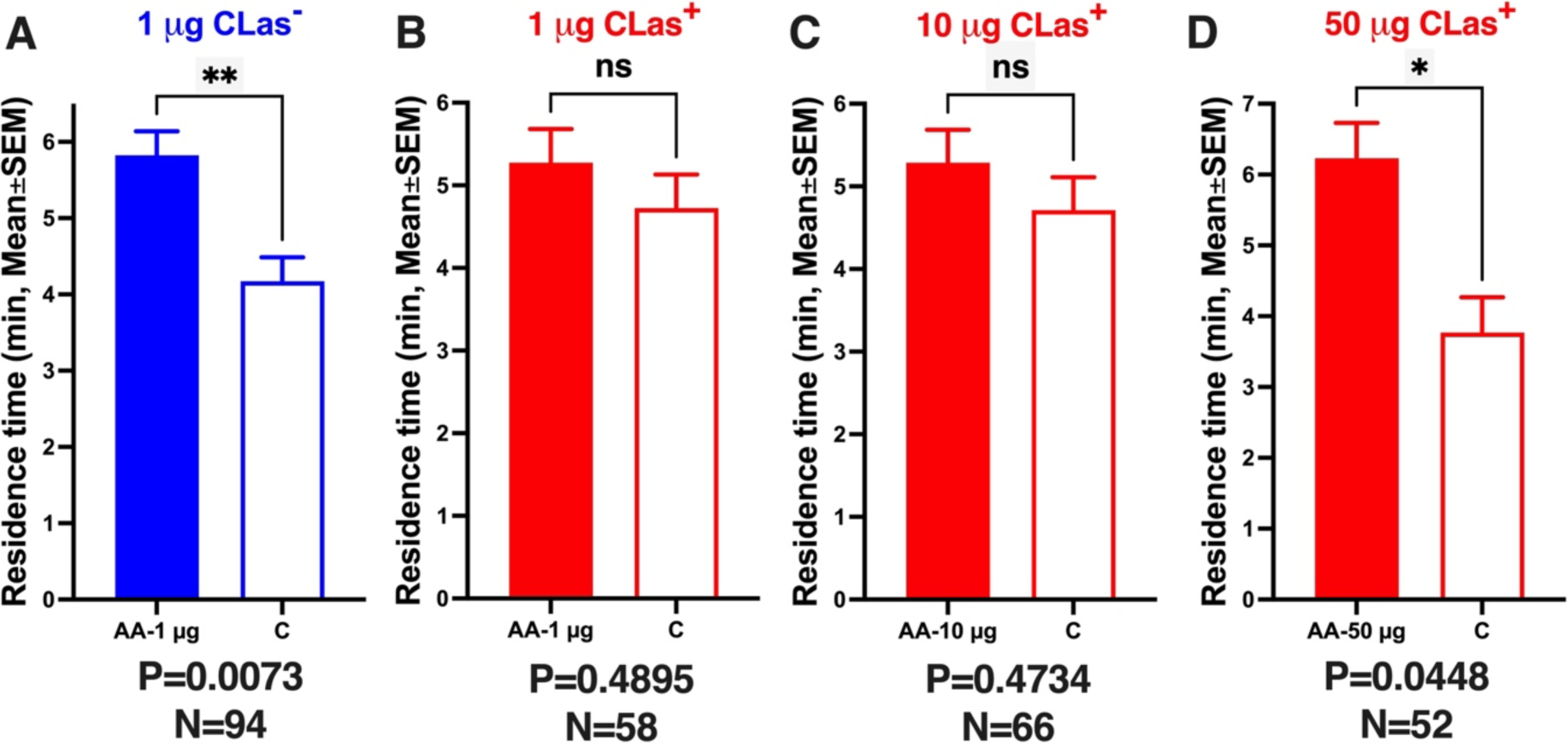
Behavioral responses from non-infected and CLas-infected ACP males to acetic acid. (A) CLas-free ACP males were tested at 1 µg (source dose). (B), (C) and (D) CLas+ ACP males at 1, 10, and 50 µg doses, respectively. Bars represent mean residence times in each odorant field ± SEM. N represents the number of male responders. qPCR analyses were used to select only CLas+ males for analysis groups (B, C, and D). We used untransformed data for normality and significant tests. P˂0.05 denotes a significant difference in the control and treatment odor fields (Wilcoxon test).

Previously, we have demonstrated that acetic acid is attractive at 1 µg, but not at 0.1 or 10 µg^20, 21^. We recapitulated these findings by testing acetic acid at 10 µg. In agreement with our previous behavioral measurements,^2, 19^ there was no significant difference between male responses to acetic acid (10 µg source dose) or control. Specifically, 74 of 112 tested males responded and spent 5.1 ± 0.3 min in the 10 µg acetic acid treatment and 4.9 ± 0.3 min in the control field of the arena (P = 0.9349, Wilcoxon test). Seven-day-old virgin females (N = 73; tested, 92) tested as a control spent 5.5 ± 0.37 min in the 10 µg acetic acid treatment field and 4.5 ± 0.37 min in the control field (P = 0.2143, Wilcoxon test). In summary, CLas-uninfected male ACP showed a significant preference for acetic acid (1 µg source dose) compared to the control (Figure 3A) but not for acetic acid at 10 µg source dose. At the higher dose (10 µg), we observed a significantly higher preference for the control field when the tested male psyllids made the first choice. They visited the control and treatment fields 0.62 ± 0.06 and 0.38 ± 0.06 times, respectively (P = 0.0474, Wilcoxon test). By contrast, these tested males significantly preferred the treatment as the final choice (control, 0.36 ± 0.06 visits; treatment, 0.64 ± 0.06 visits; P = 0.0265, Wilcoxon test). These observations suggest that this high dose (10 µg) may cause a repulsive behavior followed by attraction once the concentration declines during behavioral measurements. We then concluded that it was unnecessary to test higher doses.

### Olfactory responses of CLas-infected (CLas^+^) ACPs to acetic acid

Having confirmed that 1 µg is the optimal dose for male attraction in our multiple-choice olfactometer, we tested CLas-infected psyllids. Unlike CLas-uninfected males, CLas+ ACP males did not prefer acetic acid at 1 µg (Figure 3B). It is worth mentioning that both CLas+ and CLas-ACPs were *Wolbachia* positive. CLas+ males spend, on average, 5.3 ± 0.41 min in the treatment field and 4.7 ± 0.41 min in the control field (N = 58, P = 0.4895, Wilcoxon test). After each experiment, we analyzed each behavioral responder by qPCR to ascertain that tested ACPs were CLas+.The behavioral data used in the analysis are from 58 confirmed CLas+ responders. CLas+ females were tested at the same dose as a negative control from a behavioral perspective. These CLas+ females showed no field preference (treatment, 5.1 ± 0.38 min; control, 4.9 ± 0.38 min; N = 77, P = 0.7015, Wilcoxon test).

Next, we measured the behavioral responses of CLas+ males to a higher dose of acetic acid (10 µg). We observed no significant difference in the residence times in the treatment and control fields (Figure 3C). CLas+ males (N = 66) spent 5.3 ± 0.40 min in the treatment field and 4.7 ± 0.40 min in the control field (P = 0.4734, Wilcoxon test). Likewise, responses from CLas+ females (N = 53) were not significantly different (P = 0.7838, Wilcoxon test; control, 5.1 ± 0.47 min; treatment, 4.9 ± 0.47 min). It did not escape our attention that CLas+ males significantly preferred the arena’s 10 µg acetic acid treatment field in their first choice (treatment 0.64 ± 0.06 visits; control 0.36 ± 0.06 visits; P = 0.0356, Wilcoxon test). We surmised that this dose was not high enough to retain activity throughout the entire duration of the experiment, thus leading to no significant difference in the overall residence times (P = 0.4734, Wilcoxon test; see above). This observation prompted us to test a higher dose (100 µg). CLas+ male mean residence times in the control and 100 µg of acetic acid fields were not significantly different (N = 47, control 4.7 ± 0.52 visits; treatment, 5.3 ± 0.52 visits; P = 0.5777, Wilcoxon test). Additionally, the first choice preference for the treatment field was not retained at this higher dose (treatment 0.49 ± 0.07 visits; control 0.51 ± 0.07 visits; N=47, P=0.9999). We concluded that this dose (100 µg of acetic acid) was too high, while the lower dose (10 µg) was too low, and speculated that an intermediate dose might attract CLas+ males. We measured CLas+ male responses to 50 µg of acetic acid to test this assumption. CLas+ males showed a significant preference for the treatment field of the olfactometer (Figure 3D; N = 52, treatment, 6.2 ± 0.5 min; control, 3.8 ± 0.5; P = 0.0448, Wilcoxon test). In summary, CLas infection affected male responses to the putative sex pheromone. Specifically, CLas-infected males required a 50x higher dose to respond to acetic acid in an olfactometer.

### Electrophysiological responses of CLas- and CLas+ to acetic acid

Lastly, we compared the electrophysiological responses of uninfected and CLas+ ACP males to acetic acid. We tested 7-day-old virgin males for consistency and compared their responses at two different source doses, reflecting the 50:1 ratio observed in behavioral experiments. We used freshly prepared samples in paraffin oil. The responses to acetic acid were corrected by subtracting the background responses to paraffin oil recorded from the same preparation. On average, puffing only paraffin oil recorded a response of 0.33 ± 0.28 mV (N = 12).

Uninfected males generated 0.93 ± 0.22 and 3.90 ± 0.72 mV (N = 6) when challenged with acetic acid (source dose in paraffin oil, 1 and 50 µg/µl, respectively). By contrast, CLas+ males gave more robust responses: 2.90 ± 0.77 and 12.81 ± 1.6 mV (N = 6) when stimulated with acetic acid at the source dose of 1 and 50 µg/µl, respectively (Figure 4). As the dataset passed the Shapiro-Wilk normality test, they were analyzed by unpaired, two-tailed t-test. The EAG responses elicited by uninfected and CLas+ ACP males were significantly different when compared for each dose: 1 µg/µl (P = 0.0351, t-test) and 50 µg/µl (P = 0.0043, t-test) (Figure 4). In summary, CLas-infected males responded significantly more to acetic acid than uninfected males.

**Figure 4.**
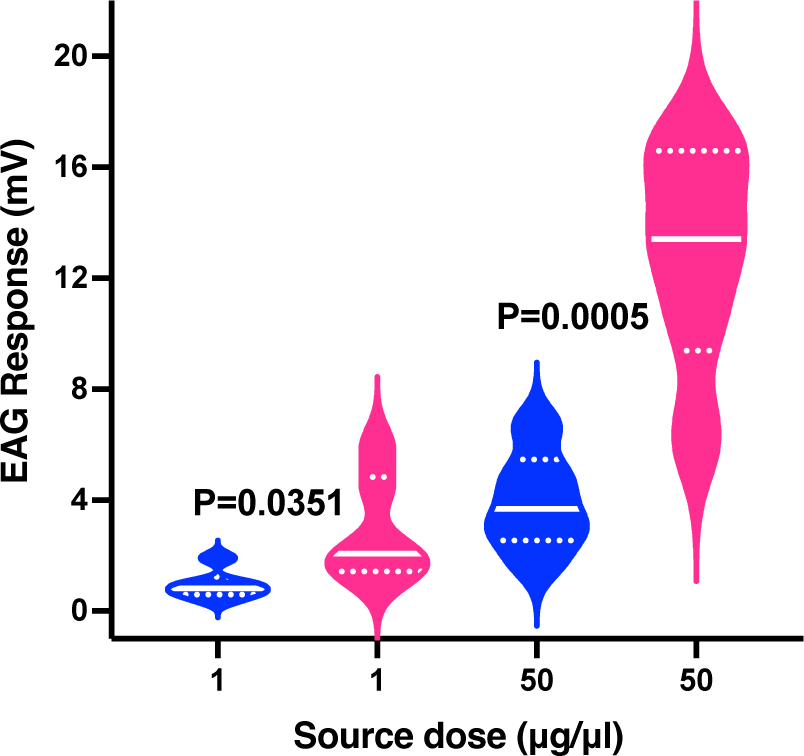
The violin plot represents the electroantennographic (EAG) responses from uninfected and CLas-infected ACP males. Statistical analyses were performed with the raw data after subtracting the background responses in each preparation to paraffin oil. The mean responses elicited by acetic acid in CLas+ males differed significantly (P<0.005, t-test) from the corresponding mean responses recorded from uninfected males at the same dose. The solid and dotted lines represent the median and quartiles in each plot. Blue plot: responses recorded with uninfected ACP males; pink plot: EAG responses elicited by CLas-infected ACP males.

### Concluding remarks

Consistent with chemical ecology terminology, we ascribed the function of acetic acid in ACP chemical communication as a “putative” sex pheromone^20^. Analytical tools did not allow us to demonstrate unequivocally that acetic acid is emitted only by ACP females. Our solvent-free, solid-phase micro-extraction (SPME) data (Figure S8 in^20^) showed that the titers of acetic acid in males and females did not differ in samples collected during no sex activity. By contrast, the titer of acetic acid in SPME samples collected from ACP females was higher than similar samples obtained from males at the same mating activity^20^. Behavioral measurements demonstrated that acetic acid is a sexual attractant, but there is no unequivocal evidence showing that this semiochemical is emitted only by ACP females. Such experimental evidence is challenging because of this semiochemical’s low molecular weight and almost ubiquitous nature. Luo and collaborators^40^ recently revisited ACP chemical communication by identifying the components of *D. citri* sex pheromones extracted by SPME and organic solvents. They concluded that acetic acid was detected only in female n-hexane extracts. This conclusion is intriguing because acetic acid is usually occluded in a gas chromatography solvent front, thus preventing identification. In their publication, a table summarizing the chemicals collected by SPME indicates the absence of acetic acid in male samples. However, they also found other intriguing “female-specific” compounds like toluene. In the nonexistence of data showing that acetic acid is emitted by ACP females while calling (and not by males), it is prudent to refer to this semiochemical in the context of ACP chemical communication as a “putative” sex pheromone or a sex attractant.

To the best of our knowledge, this is the first report of a pathogen infection affecting a vector’s response to a sex attractant. Recently, a volatile sex attractant has been identified from the tsetse fly, *Glossina morsitans*^41, 42^. Although it was also demonstrated that infection with trypanosomes alters the sexual behavior of tsetse flies, no direct evidence showed that infection modifies the response to the sex attractant.

While this paper was under review, an interesting grove-level analysis of titer and prevalence of CLas and *Wolbachia* was reported from Florida, where all citrus groves are infected with greening.^3^ The authors concluded that CLas-free ACPs tend to have higher *Wolbachia* titers than CLas-infected psyllids. These exciting findings are beyond the scope of our research. Additionally, it would be challenging, if at all possible, to test this hypothesis on the ACP population from the experimental organic plot in Mogi Mirim, where nearly 100% of the psyllids were CLas+.

Microorganisms may alter insect behaviors by affecting the peripheral olfactory and/or the central nervous systems.^43^ For example, bacteria and viruses infections may lead to increased transcript levels of the odorant receptor coreceptor,^44, 45^ Orco, or an odorant receptor,^46^ thus enhancing the sensitivity of the peripheral olfactory system. It is conceivable that a similar mechanism led to the significantly higher EAG responses recorded from CLas+ ACPs (Figure 4). Testing this hypothesis must wait for the identification of the acetic acid-detecting ionotropic or odorant receptors. Microorganisms infections may also negatively affect insect behavior through the central nervous system, as reported for the West Nile virus manipulating *Culex* mosquito host-seeking behavior.^47^ As CLas infects multiple ACP tissues,^48^ it is plausible that this bacterium may positively affect the peripheral nervous system by increasing transcript levels of acetic acid-detecting receptors while negatively affecting behavioral responses to acetic acid by infections to the central nervous system.

From the perspective of vector biology, discovering the altered behavior opened new research avenues to address if/how reducing the sensitivity to a putative sex pheromone may benefit the bacterium. From the monitoring perspective, CLas infection’s effect on ACP behavior is a significant setback. It would be challenging to industrialize non-generic pheromone lures. As HLB-infected and healthy insects respond differently to acetic acid, acetic acid-based lures must be tailored for monitoring the ACP populations in infected and non-infected areas. The altered sensitivity to acetic acid would probably not be a problem in mating disruption. After all, the foundation of mating disruption is to permeate the air with pheromone concentrations above those produced by the target insect^19^. Thus, in principle, a high dose of acetic acid may lead to mating disruption of healthy and HLB-infected psyllids.

In contrast to chemical treatments (which target a weak link common to all insect species, such as blocking a sodium channel inhibiting acetylcholinesterase), chemical ecology is generally species-specific. This specificity is priceless from an environmental perspective but costly for practical applications as it requires in-depth fundamental research. The present study suggests that we still need to gain an in-depth knowledge of ACP biology to provide chemical ecology-based alternative means to monitor and control populations of the vector of this devastating disease.

## Materials and Methods

### Field tests

Two experiments were carried out simultaneously in an organic experimental ‘Tahiti’ lime (*Citrus* × *latifolia*) orchard located in Mogi Mirim, São Paulo, Brazil, with a spacing of 7.5 × 3.5 m (row and plants spacing, respectively) from October 25 to November 1, 2019. Yellow sticky cards (30 cm in length × 10 cm in width) with a central hole (2 cm in diameter) were used to assess the attractiveness of the lures.

In a plot, we compared traps loaded with acetic acid in homemade slow-release devices (ethylene-propylene side-by-side fibers-ES lures, Chiso Co. Ltd, Japan) with yellow sticky cards (without attractive compound; control traps). One hundred microliters of an acetic acid solution in hexane (0.01µg/µL) were loaded into each ES fiber once a day.

In a second plot, we compared captures in traps with a long-lasting, slow-releasing formulation and those in control traps (without attractive compounds). The long-lasting formulation consisted of a brown polyethylene bag of 5.5 cm width x 3.5 cm length manufactured by ChemTica International S. A. (Santo Domingo, Costa Rica). For each plot, we used 15 control and 15 treatment traps arranged in two rows (15 m apart) with an intertrap distance of 10.5 m (in the same row). ACP male captures were recorded daily for eight days. The number of males per trap per day was used to perform the analyses. Field tests with double comparisons (Control × ES lures – plot 1 and control × ChemTica-A – plot 2) were analyzed by Mann-Whitney test (P<0.05) with Prism 8 (GrapPad, La Jolla, CA).

### Insect preparations, CLas-negative ACP rearing

The Asian Citrus Psyllid used in this study derived from a colony kept at Fundecitrus, reared on healthy CLas-negative orange jasmine plants (common Brazilian name “murta”), *Murraya paniculata* (L.) Jack (Sapindales: Rutaceae)^49^. Briefly, orange jasmine plants were pruned to 25-30 cm tall. Soon after new shoots appeared, approximately 7 to 12 days later, eight plants were caged in 60 x 60 x 60 cm mesh boxes and placed in a greenhouse. Each cage housed 400 adult psyllids (20-day-old mated males and females) for seven days to allow oviposition. Adults were removed, and cages (housing seedlings with eggs) were maintained in the same greenhouse until nymphs reached the fifth instar. At that point, cages were transferred to a climate-controlled room at 25 ± 2 °C, 65+10 % relative humidity (RH), under a photo regime of light/dark, 14:10 hours, and luminosity of 3,000 lux. Newly emerged adults (until 24 h-old) were collected daily from the rearing cages, sexed, and confined in new orange jasmine plants (two separated groups of plants – for male and female) to guarantee age control and virgin status condition. Seven-day-old virgins, males and females, were used in indoor behavioral assays.

### CLas-positive ACP rearing

CLas-infected psyllids were raised in a separate climate-controlled room following the same rearing protocol, temperature, RH, photoregime and light intensity described above. A group of CLas-infected ‘Valencia’ sweet orange plants (*Citrus* x *sinensis* (L.) Osbeck) grafted onto ‘Rangpur’ lime rootstock (*Citrus limonia* (L.) Osbeck), all growing in a greenhouse on 2.5 L citrus pot filled with a substrate containing a sterilized mixture composed of 80% *Pinus* sp. bark, 15% vermiculite, and 5% charcoal (Multiplant Citrus; Terra do Paraíso, Holambra, SP, Brazil) and regularly irrigated with water and weekly fertigated with a solution of minerals were used as host plants. The original budstick source of CLas for the inoculation of the plants came from the Brazilian strain 9PA^50^. Before use for ACP rearing, sweet orange plants (1.5 years after bud inoculation) were tested by qPCR for CLas titer. Plants with more than 5.19 bacterial cells per gram of tissue^51^ were used. CLas citrus plants were transferred to a climate-controlled room (room conditions described above) and pruned to 50 cm in height to stimulate the production of young shoots. Eight citrus plants with V2 flush stage^52^ were caged in 60 x 60 x 60 cm mesh boxes and infested with healthy CLas-negative mated females (3 females per citrus flush) for 7 days to allow oviposition. Adults were removed, and cages (housing seedlings with eggs) were maintained in the same climate-controlled room until adult emergence. Daily, newly emerged adults (until 24 h-old) – hereafter referred to as adults F0 were collected from the rearing cages, sexed, and confined in orange jasmine plants (separated group of plants – for male and female) to guarantee age control and virgin status condition. Seven-day-old virgins, males and females, were used in indoor behavioral assays.

### ACP samples for Wolbachia or CLas detection

Insects collected in the field and tested indoor behavioral assays were stored individually in 1.5-mL microtubes and kept at −20°C until DNA extraction within two months.

### Extraction of total DNA from ACPs

The total DNA was extracted from the entire body (head + thorax + abdomen) of a single ACP sample. Firstly, the frozen ACPs were disrupted on *Tissuelyzer* equipment (Qiagen) by a metallic bead (2 mm in diameter). The total DNA was extracted using the CTAB method (cetyltrimethylammonium bromide buffer) following a previously published protocol^53^. After disruption, each sample received 1 mL of CTAB buffer (added 0.2% of β-mercaptoethanol) and kept in a water bath for 30 min at 65°C, then added 0.5 mL of chloroform: isoamyl alcohol 24:1 (v:v) and centrifuged (12.000 rpm, 5 min, 24°C), 0.3 mL of supernatant were recovered and transferred to a new 1.5-mL microtube containing 0.18 mL of isopropanol alcohol. After 30 min storage at −20°C, the samples were centrifuged (12.000 rpm, 20 min, −4°C), and the pellet was washed twice with 70% ethanol, followed by centrifuging (12.000 rpm, 10 min, −4°C). Finally, DNAs were eluted in 30 μL of Milli-Q filtered water and stored at –20°C until subsequent analysis.

### Quantitative polymerase chain reaction (qPCR) procedure

All qPCR reactions were performed in a StepOne Plus thermocycler (Applied Biosystems), with primers and probes from Macrogen, Seul, South Korea. CLas detection in ACPs was done individually using a hydrolysis probe-based qPCR system (Path-ID master mix), detecting 16S rDNA region using the following probe-primers combination^54^: probe HLBp FAM), 5’ FAM-AGACGGGTGAGTAACGCG-3’BHQ-1; forward primer HLBas, 5’-TCGAGCGCGTATGCAA-TACG-3’, and reverse primer HLBr, 5’-CTACCTTTTTCTACGGGATAACGC-3’. An additional primer-probe set based on the *D. citri* wingless gene^55^ was used as a positive internal control to determine the quality of the extracted DNA samples. This set contained the probe DCP 5’HEX-TGTGGGCGAGGCTACAGAAC-3’BHQ-1; forward primer DCF, 5’-TGGTGAAGATGGTTGTGATCTGATGTG-3’ and reverse primer DCR, 5’-AGTGGCAGCACCTTGCCA-3’. The reaction Mix (final volume:12 µL) had 0.5 µM of each 16S rDNA forward and reverse primer, 0.2 µM of DCP primers and DCp probe (0.35 μM), 6.0 µL of TaqMan qPCR Master Mix (Ambion/ThermoFisher Scientific), and 3.0 µL of total DNA. For 16S rDNA and *D. citri,* wingless gene cycling parameters were 50°C for 2 min, 95°C for 10 min, followed by 45 cycles of 95°C for 15 s and 58°C for 45 s. *Wolbachia* presence was detected by a qPCR test using the primer-pair^56^ ftsZ-F 5’-AGCAGCCAGAGAAGCAAGAG-3’ and ftsZ-R 5’-TACGTCGCACACCTTCAAAA-3’ using 1 μL of DNA and the following cycling conditions 50°C for 2 min, 95°C for 10 min, followed by 45 cycles of 95°C for 15 s and 58°C for 30 s, followed by a melting curve from 60 to 95 °C using SYBR Green PCR Master Mix. Non-template controls were used for each run.

### Electroantennogram (EAG) recording

The EAG protocol was similar to a previously described method^20, 21^. Specifically, an ACP virgin male (7-day old) was inserted into a disposable pipette tip to immobilize the body under a stereoscopic microscope (SZT, BEL Engineering, MB, Italy), except for the head that protruded from the tip. A cotton plug was inserted into the pipette’s posterior end to prevent it from crawling backward. Electric contact was achieved by using 0.39 mm gold wires (Sigmund Cohn Corp, Mt. Vernon, NY) inserted into glass capillaries filled with Ringer’s solution (saline solution 3.7 g NaCl, 0.175 g KCl, 0.17 g CaCl_2_ in 500 mL of distilled water). The reference electrode was inserted into the head, and the recording electrode was in contact with the antennal tip. The stimulus delivery system employed was loaded on a filter paper stripe (1 × 1 cm^2^) in a disposable glass Pasteur pipette cartridge. The stimuli were delivered over the preparation in a constant 1 L/min airstream (CS-55 Stimulus Controller, Syntech) and applied (2 s duration) every 30 s interval. A 10- µl aliquot of acetic acid solution (1 µg/µL or 50 µg/µL diluted in paraffin oil – Sigma Aldrich, Milwaukee, WI) was applied to strips of filter paper and placed into the cartridge. Tested doses were alternated with paraffin oil controls to allow for the decline in the EAG response of the preparations with time. Each dose and control were tested only once for an antenna (repetition). The recorded signal was amplified using a pre-amplification probe (Universal AC/DC probe, Syntech, Germany), which was connected to an EAG high-impedance amplifier (IDAC-2, Syntech, Germany). The digitized signals were processed with EAG-Pro (version 2.0, Syntech, Germany) and were computed as the difference between the baseline and the maximum amplitude reached during odor stimulation. Antennae from six CLas-infected and six healthy ACP males were tested. The CLas-infected ACP males were subsequently analyzed by qPCR to confirm they were indeed CLas+. For each AA concentration repetition (10 µg and 500 µg), the relative EAG response was calculated by subtracting the EAG response (mV) value of the paraffin oil (control) from the EAG response value for the AA doses related to the same antennae.

### Indoor bioassays

All behavioral assays were done in a climate-controlled room at 25 ± 2 °C, 65 ± 10% relative humidity, 14 h light/10 h dark photo regime, and 3,000 lux luminosity. Responses to acetic acid were measured with a previously described multi-choice olfactometer^20, 39^. In brief, we used an acrylic 4-arm olfactometer (30.0 × 30.0 × 2.5 cm; length × width × height, respectively) with a transparent acrylic lid and modified by adding a yellow background below the bottom of the device ^39^. Compressed air (charcoal filtered and humidified) was connected to a stainless-steel line and split into four individual 0.635-cm-diameter polytetrafluoroethylene (PTFE) tubes (Sigma-Aldrich, Bellefonte, PA, USA) connected in four flowmeters [0.1–1 LPM, Brooks Instruments, Hatfield, PA, USA], adjusted to 0.1 LPM airflow/flowmeter. Each PTFE tube was connected to one horizontal glass chamber (20 cm length × 6 cm internal diameter) containing an odor source (two sources of acetic acid, treatment and 2 of hexane, control; both loaded on cotton swabs), and each airflow converged through PTFE tubes to one of the four device arms.

### Statement for guidelines and permission

The FUNDECITRUS (Fund for Citrus Protection) is a non-profit association maintained by citrus growers and juice manufacturers from the State of São Paulo to foster the sustainable development of the citrus industry. The FUNDECITRUS’s research activities focus mainly on managing citrus pests and diseases using citrus growerś financial support. As part of this agreement, the growers allow the FUNDECITRUS free access to their citrus orchards to develop research that benefits the management of citrus pests and diseases. Therefore, all fieldwork in São Paulo is appropriately authorized and conforms to institutional guidelines. Additionally, the FUNDECITRUS uses its own facility to grow infected and non-infected plant varieties and raises insects for research.

## Supporting information

Dataset with raw data for figures

## Acknowledgments.

We thank Jean M. Martins and Deividson F. Rodrigues (FUNDECITRUS) for DNA extractions and Daniela A. B. Coletti (FUNDECITRUS) for performing qPCR reactions, Drs. Rodrigo Facchini Magnani (FUNDECITRUS) and Francisco Gonzales (ChemTica International) for critically reading an earlier manuscript draft, and ChemTica International for providing experimental acetic acid-based lures. This work was supported partially by the Fund for Citrus Protection (FUNDECITRUS) under a research agreement with the University of California-Davis (#201600147) and the National Institute of Science and Technology of Semiochemicals in Agriculture (INCT) [grants FAPESP #2014/50871-0 and CNPq #465511/2014-7].

## Author contributrions

W.S.L. and H.X.L.V. designed research. H.X.L.V., M.C.S., R.A.G.L., R.F., V. E., J.C.D., A.A.L.P., D.M.M., A.P.F. performed research. N.A.W., M.P.M., and J.M.S.B. provided reagents, tools, and ideas. W.S.L. wrote the manuscript. All authors reviewed and approved the final version of the manuscript.

## Additional Information

A dataset with raw data used to generate figures accompanies this paper at xxxx

## Data Availability

All data generated during this study are included in the manuscript and supporting files.

