## Supplementary material for "The greening-causing agent alters the behavioral and electrophysiological responses of the Asian citrus psyllid to a putative sex pheromone": Dataset with raw data for figures

**Raw data for Figure 1 (2 pages)**

| 2/18 | 2/20 | 3/9 | 3/13 | 3/16 | 29/04 | 5/4 | 5/9 | 5/13 | 5/18 | 5/23 | 6/2 | 6/6 |
|---|---|---|---|---|---|---|---|---|---|---|---|---|
| 1 | 0.54 | 1 | 1.5 | 1.22 | 0.82 | 0.54 | 0.67 | 1.22 | 0.54 | 0.67 | 0.33 | 0.54 |
| 0.43 | 0.4 | 1.5 | 0.67 | 0.43 | 0.82 | 0.33 | 0.67 | 0.25 | 0.67 | 0.43 | 0.54 | 0.54 |
| 1 | 0.54 | 1.22 | 0.82 | 0.67 | 1.5 | 1.5 | 0.33 | 0.33 | 0.67 | 1.22 | 0.67 | 0.67 |
| 0.11 | 1 | 1.22 | 0.54 | 0.82 | 0.33 | 0.67 | 0.54 | 0.67 | 0.82 | 0.43 | 0.67 | 0.54 |
| 0.38 | 0.5 | 0.54 | 0.54 | 0.43 | 0.82 | 1 | 0.54 | 0.54 | 1.5 | 1.22 | 0.31 | 0.67 |

| 6/13 | 6/15 | 6/26 | 07/07 | 7/10 | 8/28 | 9/18 | 9/25 | 10/9 | 10/22 | 11/27 | 12/22 |
|---|---|---|---|---|---|---|---|---|---|---|---|
| 0.43 | 0.67 | 0.33 | 0.54 | 0.33 | 1.1 | 0.67 | 0.82 | 0.43 | 0.67 | 1.5 | 0.82 |
| 0.43 | 0.33 | 0.33 | 1 | 0.18 | 0.54 | 0.82 | 1.5 | 1.5 | 1.22 | 1.22 | 0.33 |
| 0.33 | 0.54 | 0.25 | 0.67 | 0.25 | 0.31 | 0.43 | 0.43 | 0.67 | 1.22 | 1.22 | 0.18 |
| 0.33 | 0.82 | 0.54 | 1 | 0.33 | 0.31 | 0.25 | 0.67 | 0.5 | 1 | 0.67 | 1 |
| 0.18 | 0.67 | 0.25 | 0.18 | 0.43 | 1 | 0.67 | 0.82 | 1.22 | 0.54 | 1.22 | 0.67 |

**Raw data for Figure 2**

| Male | Female | Male | Female |
|---|---|---|---|
| 96.77 | 100 | 100 | 100 |
| 100 | 100 | 100 | 100 |
| 100 | 97.96 | 100 | 100 |
| 100 | 100 | 100 | 98.25 |
| 100 | 100 | 97.44 | 100 |
| 100 | 100 | 93 | 100 |
| 100 | 100 | 100 | 94.9 |
| 100 | 100 | 100 | 100 |
| 100 | 100 | 90.8 | 97.1 |
| 100 | 100 | 95.7 | 100 |
| 100 | 98.2 | 98.2 | 100 |

**Raw data for Figure 3 (3 pages)**

| **Figure 3A** | | **Figure 3B** | | **Figure 3C** | | **Figure 3D** | |
|---|---|---|---|---|---|---|---|
| **AA-1 µg** | **C** | **AA-1 µg** | **C** | **AA-10 µg** | **C** | **AA-50 µg** | **C** |
| 10 | 0 | 5.56 | 4.44 | 5.08 | 4.92 | 10 | 0 |
| 8.1 | 1.9 | 4.1 | 5.9 | 6.97 | 3.03 | 10 | 0 |
| 1.99 | 8.01 | 10 | 0 | 9.46 | 0.54 | 0 | 10 |
| 4.13 | 5.87 | 2.05 | 7.95 | 0 | 10 | 10 | 0 |
| 8.54 | 1.46 | 6.2 | 3.8 | 6.83 | 3.17 | 6.89 | 3.11 |
| 10 | 0 | 4.28 | 5.72 | 4 | 6 | 0 | 10 |
| 0 | 10 | 6.06 | 3.94 | 4.47 | 5.53 | 9.09 | 0.91 |
| 4.14 | 5.86 | 0 | 10 | 1.86 | 8.14 | 7.34 | 2.66 |
| 6.57 | 3.43 | 2.56 | 7.44 | 8.16 | 1.84 | 0 | 10 |
| 1.94 | 8.06 | 0 | 10 | 10 | 0 | 6.46 | 3.54 |
| 1.66 | 8.34 | 0 | 10 | 10 | 0 | 1.1 | 8.9 |
| 3.88 | 6.12 | 5.98 | 4.02 | 6.88 | 3.12 | 8.05 | 1.95 |
| 1.32 | 8.68 | 5.45 | 4.55 | 2.87 | 7.13 | 9.78 | 0.22 |
| 7.29 | 2.71 | 9.12 | 0.88 | 5.18 | 4.82 | 8.27 | 1.73 |
| 6.29 | 3.71 | 7.53 | 2.47 | 1.8 | 8.2 | 8.27 | 1.73 |
| 6.9 | 3.1 | 9.76 | 0.24 | 9.39 | 0.61 | 6.01 | 3.99 |
| 10 | 0 | 10 | 0 | 6.45 | 3.55 | 7.7 | 2.3 |
| 8.33 | 1.67 | 10 | 0 | 7.12 | 2.88 | 4 | 6 |
| 3.87 | 6.13 | 5.17 | 4.83 | 0 | 10 | 9.65 | 0.35 |
| 8.12 | 1.88 | 6.65 | 3.35 | 1.56 | 8.44 | 7.06 | 2.94 |
| 6.44 | 3.56 | 7.06 | 2.94 | 8.84 | 1.16 | 5.12 | 4.88 |
| 0 | 10 | 0.97 | 9.03 | 10 | 0 | 0 | 10 |
| 6.71 | 3.29 | 6 | 4 | 3.1 | 6.9 | 8.91 | 1.09 |
| 0 | 10 | 10 | 0 | 0.34 | 9.66 | 7.68 | 2.32 |
| 9.43 | 0.57 | 8.24 | 1.76 | 10 | 0 | 10 | 0 |
| 0 | 10 | 3.21 | 6.79 | 0 | 10 | 7.89 | 2.11 |
| 7.04 | 2.96 | 4.54 | 5.46 | 10 | 0 | 3.07 | 6.93 |
| 10 | 0 | 5.89 | 4.11 | 10 | 0 | 0 | 10 |
| 8.03 | 1.97 | 4.54 | 5.46 | 0.9 | 9.1 | 7.75 | 2.25 |
| 7.87 | 2.13 | 3.73 | 6.27 | 2.34 | 7.66 | 10 | 0 |
| 8.62 | 1.38 | 4.48 | 5.52 | 6.6 | 3.4 | 3.85 | 6.15 |
| 3.3 | 6.7 | 1.81 | 8.19 | 8.02 | 1.98 | 5.82 | 4.18 |
| 8.99 | 1.01 | 8.2 | 1.8 | 3.97 | 6.03 | 0 | 10 |
| 7 | 3 | 2.71 | 7.29 | 5.4 | 4.6 | 10 | 0 |
| 6.28 | 3.72 | 8.19 | 1.81 | 6.88 | 3.12 | 1.64 | 8.36 |
| 10 | 0 | 3.89 | 6.11 | 2.82 | 7.18 | 10 | 0 |
| 6.93 | 3.07 | 9.09 | 0.91 | 7.06 | 2.94 | 9.54 | 0.46 |
| 9.91 | 0.09 | 10 | 0 | 5.87 | 4.13 | 10 | 0 |
| 7.37 | 2.63 | 5.66 | 4.34 | 6.2 | 3.8 | 2.29 | 7.71 |
| 5.3 | 4.7 | 7.08 | 2.92 | 1.12 | 8.88 | 5.85 | 4.15 |
| 0 | 10 | 0.95 | 9.05 | 7.45 | 2.55 | 0 | 10 |
| 7.85 | 2.15 | 3.53 | 6.47 | 8.86 | 1.14 | 6.06 | 3.94 |

| | | | | | | | |
|---|---|---|---|---|---|---|---|
| 5.04 | 4.96 | 0 | 10 | 6.14 | 3.86 | 8.07 | 1.93 |
| 2.29 | 7.71 | 2.03 | 7.97 | 1.03 | 8.97 | 8.81 | 1.19 |
| 4.01 | 5.99 | 4.3 | 5.7 | 6.02 | 3.98 | 6.62 | 3.38 |
| 7.82 | 2.18 | 4.3 | 5.7 | 2.07 | 7.93 | 10 | 0 |
| 2.8 | 7.2 | 4.01 | 5.99 | 1.32 | 8.68 | 9.72 | 0.28 |
| 9.3 | 0.7 | 6.16 | 3.84 | 4.02 | 5.98 | 8.63 | 1.37 |
| 4.31 | 5.69 | 2.12 | 7.88 | 8.19 | 1.81 | 6.65 | 3.35 |
| 3.96 | 6.04 | 10 | 0 | 4.05 | 5.95 | 1.27 | 8.73 |
| 9.72 | 0.28 | 3.96 | 6.04 | 6.06 | 3.94 | 9.07 | 0.93 |
| 8.22 | 1.78 | 9.01 | 0.99 | 1.22 | 8.78 | 0 | 10 |
| 10 | 0 | 6.83 | 3.17 | 10 | 0 | | |
| 0 | 10 | 0 | 10 | 10 | 0 | | |
| 6.22 | 3.78 | 0 | 10 | 10 | 0 | | |
| 9.31 | 0.69 | 6.93 | 3.07 | 2.81 | 7.19 | | |
| 5.05 | 4.95 | 6.43 | 3.57 | 3.07 | 6.93 | | |
| 7.42 | 2.58 | 9.67 | 0.33 | 1.7 | 8.3 | | |
| 6.72 | 3.28 | | | 3.3 | 6.7 | | |
| 4.4 | 5.6 | | | 6.23 | 3.77 | | |
| 6.27 | 3.73 | | | 7.56 | 2.44 | | |
| 3.78 | 6.22 | | | 9.22 | 0.78 | | |
| 0 | 10 | | | 4.66 | 5.34 | | |
| 8.97 | 1.03 | | | 2.28 | 7.72 | | |
| 0.1 | 9.9 | | | 0 | 10 | | |
| 4.96 | 5.04 | | | 4.13 | 5.87 | | |
| 8.05 | 1.95 | | | | | | |
| 2.85 | 7.15 | | | | | | |
| 3.43 | 6.57 | | | | | | |
| 4.97 | 5.03 | | | | | | |
| 6.17 | 3.83 | | | | | | |
| 4.23 | 5.77 | | | | | | |
| 10 | 0 | | | | | | |
| 8.35 | 1.65 | | | | | | |
| 4.06 | 5.94 | | | | | | |
| 5.57 | 4.43 | | | | | | |
| 3.01 | 6.99 | | | | | | |
| 7.19 | 2.81 | | | | | | |
| 8.21 | 1.79 | | | | | | |
| 7.64 | 2.36 | | | | | | |
| 10 | 0 | | | | | | |
| 7.97 | 2.03 | | | | | | |
| 5.15 | 4.85 | | | | | | |
| 6.99 | 3.01 | | | | | | |
| 2.07 | 7.93 | | | | | | |
| 1.22 | 8.78 | | | | | | |
| 8.36 | 1.64 | | | | | | |

| | |
|---|---|
| 8.36 | 1.64 |
| 0 | 10 |
| 3.02 | 6.98 |
| 8.75 | 1.25 |
| 7.37 | 2.63 |
| 6.04 | 3.96 |
| 7.77 | 2.23 |

**Raw data for Figure 4**

| Clas-free | Clas+ | Clas-free | Clas+ |
|---|---|---|---|
| **1** | **1** | **50** | **50** |
| 0.931 | 1.48 | 1.74 | 6.256 |
| 0.382 | 5.981 | 6.668 | 13.946 |
| 1.907 | 2.121 | 3.189 | 16.8 |
| 0.687 | 4.456 | 4.166 | 16.525 |
| 0.656 | 2.014 | 2.793 | 10.437 |
| 1.007 | 1.267 | 5.051 | 12.894 |
